# Visual adaptation stronger at horizontal than vertical meridian: Linking performance with V1 cortical surface area

**DOI:** 10.1101/2025.03.07.642102

**Authors:** Hsing-Hao Lee, Marisa Carrasco

**Affiliations:** Department of Psychology, New York University, New York, NY, USA; Center for Neural Sciences, New York University, New York, NY, USA

**Keywords:** Visual adaptation, Contrast sensitivity, Polar angle asymmetries, V1 surface area

## Abstract

Visual adaptation reduces bioenergetic expenditure by decreasing sensitivity to repetitive and similar stimuli. In human adults, visual performance varies systematically around polar angle for many visual dimensions and tasks: Performance is superior along the horizontal than the vertical meridian (horizontal-vertical anisotropy, HVA), and the lower than upper vertical meridian (vertical meridian asymmetry, VMA). These asymmetries are resistant to spatial and temporal attention. However, it remains unknown whether visual adaptation differs around polar angle. Here, we investigated how adaptation influences contrast sensitivity at the fovea and perifovea across the four cardinal meridian locations, for both horizontal and vertical stimuli in an orientation discrimination task. In the non-adapted conditions, the HVA was more pronounced for horizontal than vertical stimuli. For both orientations, adaptation was stronger along the horizontal than vertical meridian, exceeding foveal adaptation. Additionally, perifoveal adaptation effects positively correlated with individual V1 cortical surface area. These findings reveal that visual adaptation mitigates the HVA in contrast sensitivity, fostering perceptual uniformity around the visual field while conserving bioenergetic resources.

## Significance Statement

Human visual perception varies around the visual field, with robust horizontal-vertical and vertical meridian asymmetries. These asymmetries are pervasive –present for monocular and binocular viewing, different stimulus sizes, orientations, eccentricities, luminance levels– and resilient –they remain under conditions that improve perception, such as covert attention, and are even exacerbated with presaccadic attention. Here, we investigated whether visual adaptation, which helps manage bioenergetic resources, alters these polar angle asymmetries. We found that adaptation decreases the horizontal-vertical asymmetry, making perception more uniform across the visual field. Moreover, the degree of this effect correlates with surface area in primary visual cortex. This is the first study revealing that a fundamental visual process –adaptation– diminishes this prominent perceptual asymmetry.

## Introduction

Visual performance exhibits systematic spatial variations across the visual field: Sensitivity declines with eccentricity (1–3) and varies systematically around the polar angle: Performance is superior along the horizontal than the vertical meridian *(horizontal-vertical anisotropy,* HVA), and along the lower than the upper vertical meridian (*vertical meridian asymmetry,* VMA). These asymmetries persist across multiple dimensions —e.g., contrast sensitivity (4–10), spatial resolution (11–15), motion (16–20), visual acuity (21, 22), perceived contrast (9), perceived spatial frequency (23)— and mid-level and higher-order tasks, e.g., crowding (15, 24–27), texture segmentation (14, 15, 21, 22, 28), perceived object size (29), face perception (30, 31), short-term memory (23), and word identification (32). However, contrast sensitivity is the currency of the visual system, which most – if not all – visual dimensions depend upon to some degree.

Visual adaptation decreases sensitivity for stimuli that have been encountered repeatedly in the past, thereby increasing sensitivity to changes in the environment (33). Adaptation helps conserve the brain’s limited bioenergetic resources by allocating less energy to repetitive stimuli (33–36). For instance, contrast adaptation reduces sensitivity (35–43) and neural responses (44–53), and shifts the contrast response function in V1 to recenter sensitivity away from the adaptor (54).

However, there is a spatial bias in adaptation studies: (1) Most studies have focused on horizontal meridian locations (e.g. 55, 56-58). (2) The few studies that tested other locations (e.g., intercardinal locations or vertical meridian) have not analyzed them separately (e.g. 43, 59). Thus, whether and how the adaptation effect varies around polar angle remain unknown. (3) Moreover, the magnitude of the adaptation effect across eccentricity has yielded inconsistent findings; some show similar extent of adaptation with cortically-magnified stimuli (40, 56), whereas others show parafoveal dominance (58, 59). Addressing these gaps will reveal how adaptation alters perception and conserves bioenergetic resources throughout the visual field.

Cortical magnification –the amount of cortical surface area corresponding to one degree of visual angle (mm^2^/°)– declines with eccentricity (60–64) and has been used to link perceptual performance to brain structure. Converging neural evidence demonstrates that cortical magnification limits peripheral vision: V1 surface area across eccentricity correlates with various perceptual measures, including perceived object size (65, 66), perceived angular size (67), and acuity (68, 69). Moreover, V1 surface area around polar angle also correlates with perceptual measures, including contrast sensitivity and acuity (70–72).

Here, we investigated whether adaptation decreases contrast sensitivity similarly around polar angle and at fovea using an orientation discrimination task. This task has been a proxy for contrast sensitivity because on this task performance monotonically increases with contrast (73). This task has been used this way in many studies (e.g. 6, 7, 10, 74-84), including adaptation studies (35, 42, 85). Here, we tested three hypotheses:

1. Uniform adaptation: Similar adaptation magnitude across meridians, consistent with the central role of early visual cortex in both adaptation (36, 42) and covert spatial attention (36, 84), and with the findings that attention improves performance similarly around polar angle (86–89).
2. Vertical meridian dominance: Stronger adaptation along the vertical (particularly the upper vertical meridian) than the horizontal meridian, as predicted by adaptor-target similarity (58, 59, 90) being stronger where population receptive fields (pRF) size is larger (72, 91, 92) and stimuli are less precisely encoded (93).
3. Horizontal meridian dominance: Stronger adaptation along the horizontal than the vertical meridian (particularly upper), consistent with the larger cortical surface area devoted to the horizontal than vertical meridian (63, 70–72, 91, 92). V1 neuronal density is approximately uniform across visual space (94, 95) and locations with more neurons devoted to sensory processing elicit a stronger response to the adaptor (96, 97), which amplifies the adaptation effect (54, 98–101). This prediction is consistent with the fact that adaptation increases with the strength of the response to the stimulus (102–105).

Fourteen adults participated in all three experiments (**Figure 1).** Participants performed an orientation discrimination task, as in previous studies of contrast adaptation (35, 85, 104). In Experiment 1, participants adapted to a horizontal stimulus and discriminated whether a target Gabor–presented at the same location as the adaptor among one of four perifoveal cardinal locations–was tilted clockwise or counterclockwise from horizontal (**Figure 1A**). Given that sensitivity to gratings is higher for radial than tangential orientations (106–108), we hypothesized an exacerbated HVA when the task involved horizontal stimulus orientation, which is radial at the horizontal meridian. To evaluate the influence of radial bias, in Experiment 2 participants adapted to a vertical stimulus and discriminated whether a Gabor was tilted clockwise or counterclockwise from vertical. These experiments enabled us to (1) evaluate the role of cortical surface area in adaptation and (2) compare the extent of both HVA and VMA with different stimulus orientations.

**Figure 1.**
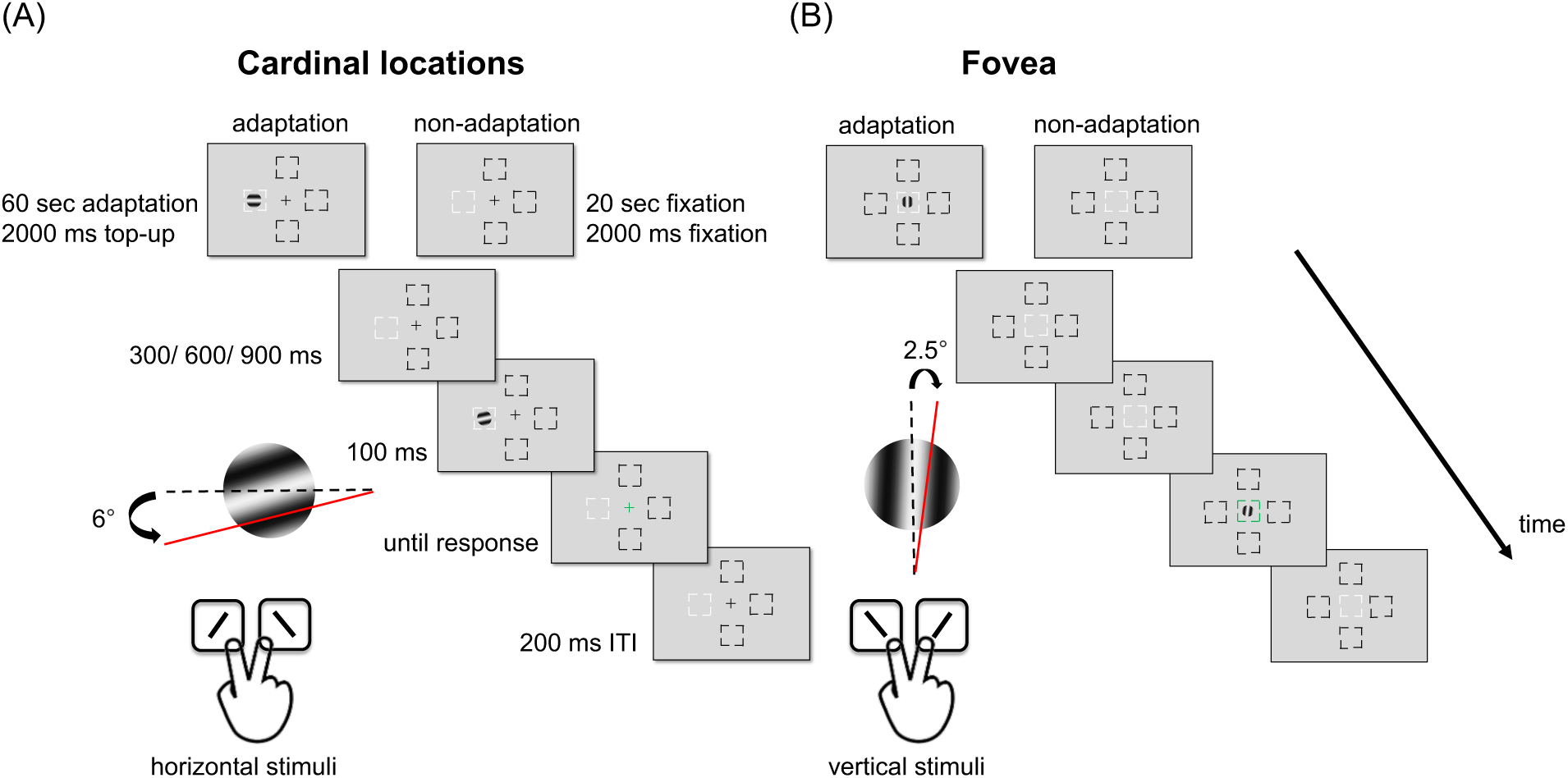
(**A**) Experimental procedure: Participants performed either adaptation or non-adaptation blocks, each in separate experimental sessions. The target Gabor stimulus was always presented within the white placeholder, and target locations were blocked. The target, a horizontal (Experiment 1) or vertical (Experiment 2) Gabor stimulus, was presented either at **(A)** the perifovea (Experiments 1 and 2) or (**B**) the fovea (Experiment 3). Participants were instructed to respond whether the Gabor was tilted clockwise or counterclockwise from horizontal (Experiment 1) or vertical (Experiment 2). The target Gabor was tilted 6° from the horizontal line or 2.5° from the vertical line. For illustration purposes, the stimulus size and spatial frequency shown here are not to scale.

In Experiment 3, we examined adaptation effects at the fovea using the same orientation discrimination tasks (**Figure 1B**). This experiment enabled us to (1) further evaluate the role of cortical surface area in adaptation and (2) address previous inconsistent findings regarding task eccentricity-dependent adaptation effects (40, 56–59, 109–111): For instance, whereas tilt aftereffects are stronger for suprathreshold targets at peripheral than central vision along the horizontal meridian (57, 58, 109–111), contrast threshold adaptation shows comparable magnitudes across foveal, parafoveal, and peripheral vision along the horizontal meridian (55), and similar recovery times for adaptation durations ≥1000 ms between peripheral and foveal vision (57).

## Results

### Experiment 1-Perifoveal Locations, Horizontal Stimulus

To investigate the adaptation effect at the vertical and horizontal meridians, we conducted a two-way ANOVA on contrast thresholds. This analysis showed a main effect of location [*F*(3,39)=14.04, *p*<.001, *η_p_^2^*=0.52] and a higher threshold in the adapted than non-adapted conditions [*F*(1,13)=45.42, *p*<.001, *η_p_^2^*=0.78], and an interaction [*F*(3,39)=4.98, *p*=.005, *η_p_^2^*=0.28], indicating that the adaptation effect varied across locations (**Figure 2A**).

**Figure 2.**
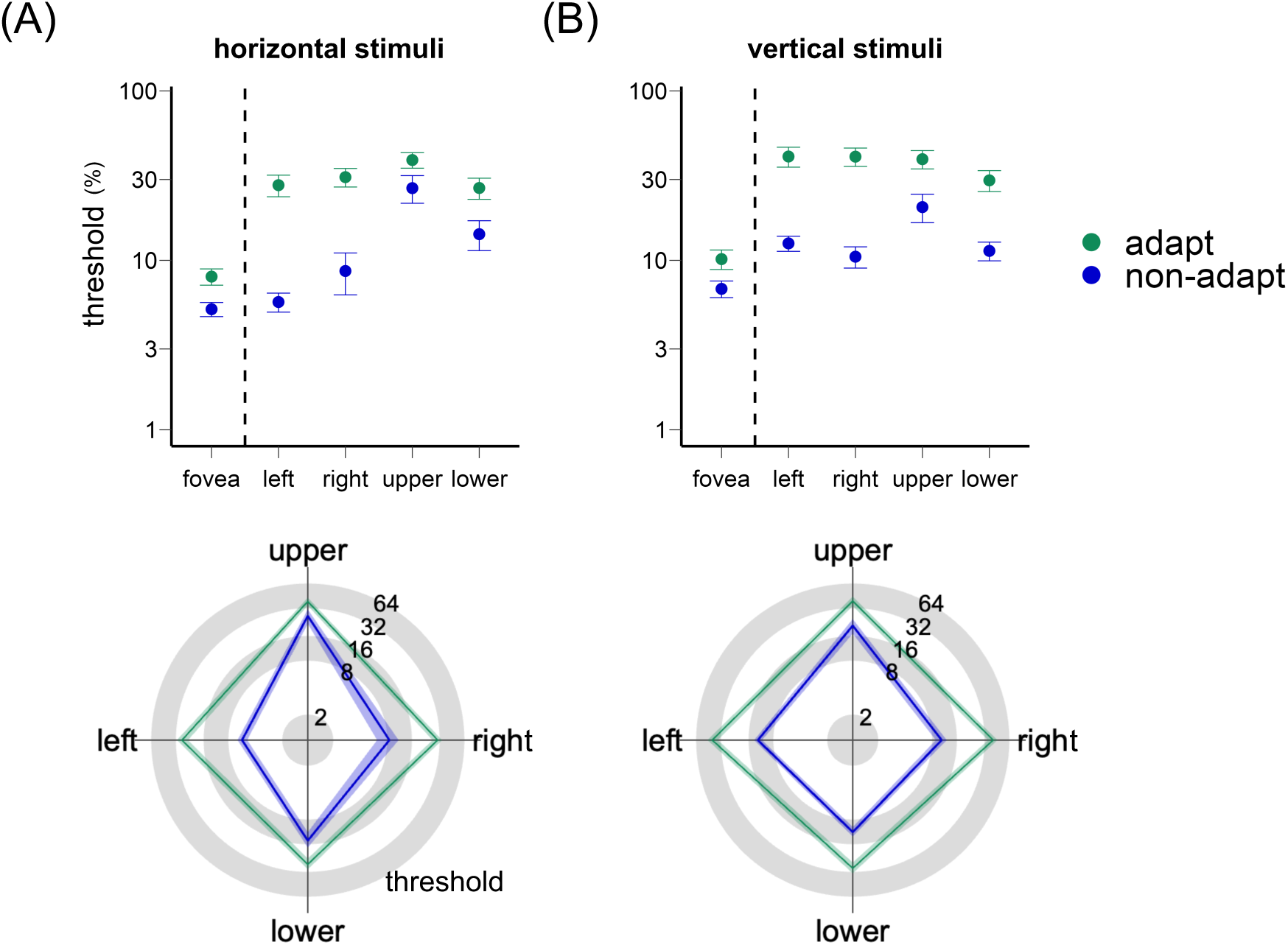
The top panel shows the contrast thresholds (log-scaled) for orientation discrimination at the fovea, and at the left, right, upper, and lower perifoveal locations for (**A**) horizontal stimuli and (**B**) vertical stimuli. Note that for the adapted condition, contrast thresholds (%) are similar for the 4 perifoveal locations (green points). The bottom panel illustrates that the adaptation effect, measured as the difference in contrast sensitivity between the adapted and non-adapted conditions, is stronger along the horizontal than the vertical meridian. The error bars represent ±1 SEM.

**Figure 3.**
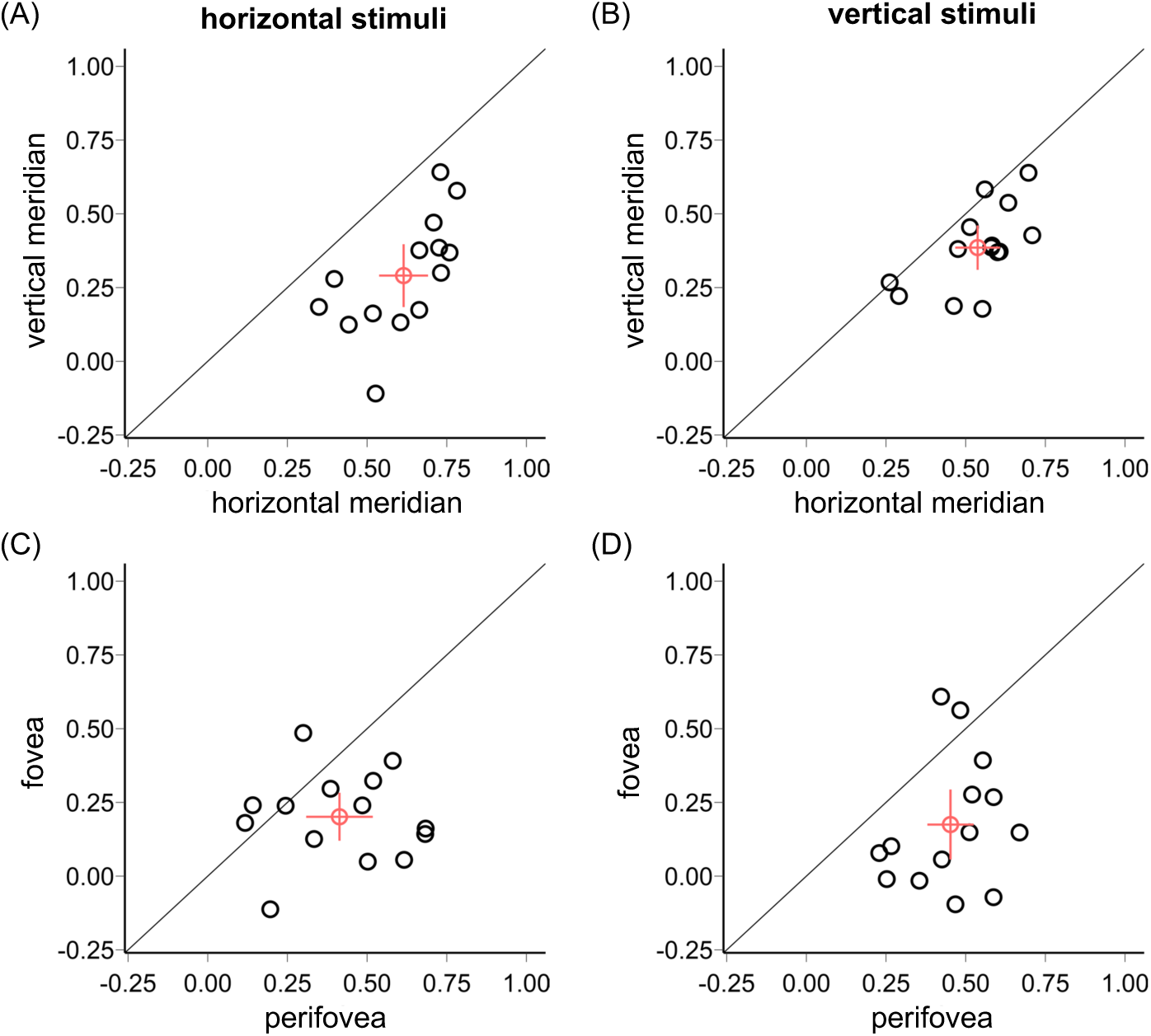
Upper panel: Normalized adaptation effects ((adapted – non-adapted threshold) / (adapted + non-adapted threshold)) are stronger along the horizontal than the vertical meridian for both (**A**) horizontal stimuli (Experiment 1) and (**B**) vertical stimuli (Experiment 2). Lower panel: Adaptation effects are stronger in the perifovea than the fovea for both (**C**) horizontal and (**D**) vertical stimuli. Each black circle represents the threshold ratio for an individual participant; the red circle indicates the mean across participants. Error bars represent ±1 SEM.

First, we confirmed the expected HVA and VMA in the non-adaptation condition (**Figure S1**, upper panel). Contrast thresholds were lower along the horizontal than the vertical meridian [*t*(13)=5.26, *p*<.001, *d*=1.41] and lower at the lower than upper vertical meridian [*t*(13)=3.15, *p*=.008, *d*=0.84].

Next, we assessed the adaptation effect at the horizontal and vertical meridians. The adaptation effect (calculated as the difference between adapted and non-adapted thresholds, as in previous studies (112–114)) was stronger at the horizontal than the vertical meridian [*t*(13)=3.77, *p*=.002, *d*=1.01] (**Figures 2A, 3A**), but there was no significant difference between the upper and lower vertical meridian [*t*(13)=0.01, *p*=.99].

To account for baseline differences in the non-adapted condition, we calculated a normalized adaptation effect [(adapted threshold–non-adapted threshold)/(adapted threshold+non-adapted threshold)]. This normalized adaption effect was also stronger at the horizontal than the vertical meridian [*t*(13)=7.84, *p*<.001, *d*=2.1], with no significant difference at the upper and lower vertical meridians [*t*(13)=1.3, *p*=.217]. In summary, the decrease in contrast sensitivity following adaptation was more pronounced at the horizontal than the vertical meridian.

### Experiment 2 – Perifoveal Locations, Vertical Stimulus

When using vertical adaptor and target stimuli, the findings were consistent with those in Experiment 1. A two-way ANOVA on contrast thresholds showed main effects of location [*F*(3,39)=4.59, *p*=.008, *η_p_^2^*=0.26] and adaptation [*F*(1,13)=44.15, *p*<.001, *η_p_^2^*=0.77], as well as an interaction [*F*(3,39)=6.63, *p*=.001, *η_p_^2^*=0.34], indicating that the adaptation effect varied across locations (**Figure 2B**).

In the non-adapted condition (**Figure S1,** lower panel), we confirmed the expected HVA and VMA. Contrast thresholds were lower along the horizontal than the vertical meridian [*t*(13)=2.18, *p*=.048, *d*=0.58] and lower at the lower than upper vertical meridian [*t*(13)=2.57, *p*=.023, *d*=0.69].

The adaptation effect (calculated as the difference between adapted and non-adapted thresholds, as in previous studies (112–114)) was stronger at the horizontal than the vertical meridian [*t*(13)=4.04, *p*=.001, *d*=1.08] (**Figures 2B, 3B**), but there was no significant difference between the upper and lower vertical meridian [*t*(13)=0.22, *p*=.831]. Similarly, the normalized adaptation effect was stronger at the horizontal than the vertical meridian [*t*(13)=4.78, *p*<.001, *d*=1.28], with no significant difference between the upper and lower vertical meridians [*t*(13)=1.54, *p*=.147].

### Comparing adaptation between stimulus orientations

A 3-way ANOVA on contrast thresholds, with factors of location, adaptation, and stimulus orientation (horizontal: Experiment 1; vertical: Experiment 2) showed main effects of adaptation [*F*(1,13)=54.22, *p*<.001, *η_p_^2^*=0.81] and location [*F*(3,39)=8.74, *p*<.001, *η_p_^2^*=0.4], but not of stimulus orientation [*F*(1,13)=1.25, *p*=.267] or 3-way interaction [*F*(3,39)<1]. All two-way interactions emerged: location x orientation [*F*(3,39)=12.33, *p*<.001, *η_p_^2^*=0.49], adaptation x orientation [*F*(1,13)=5.72, *p*=.033, *η_p_^2^*=0.31], and adaptation x location [*F*(3,39)=9.35, *p*<.001, *η_p_^2^*=0.42].

The interaction between location and orientation (across adaptation conditions) showed a stronger HVA for horizontal than vertical stimuli [*t*(13)=4.89, *p*<.001, *d*=1.31] but no difference for the VMA [*t*(13)=1.39, *p*=.187]. The interaction between adaptation and orientation (across locations) yielded a stronger adaptation effect for the vertical than horizontal stimuli [*t*(13)=2.39, *p*=.033, *d*=0.64], but this difference was not significant for the normalized adaptation effect [*t*(13)=0.78, *p*=.449]. The interaction between adaptation and location (across orientations) reflected a stronger adaptation effect for horizontal than vertical locations, both without normalization [*t*(13)=5.07, *p*<.001, *d*=1.36], and with normalization [*t*(13)=7.18, *p* <.001, *d*=1.92].

The differences in HVA and VMA between horizontal and vertical stimuli under non-adapted conditions (**Figure S1**) resulted from a stronger HVA for horizontal stimuli [*t*(13)=3.83, *p*=.002, *d*=1.03, **Figure 4A**], with no significant difference for the VMA [*t*(13)=1.04, *p*=.318, **Figure 4B**].

**Figure 4.**
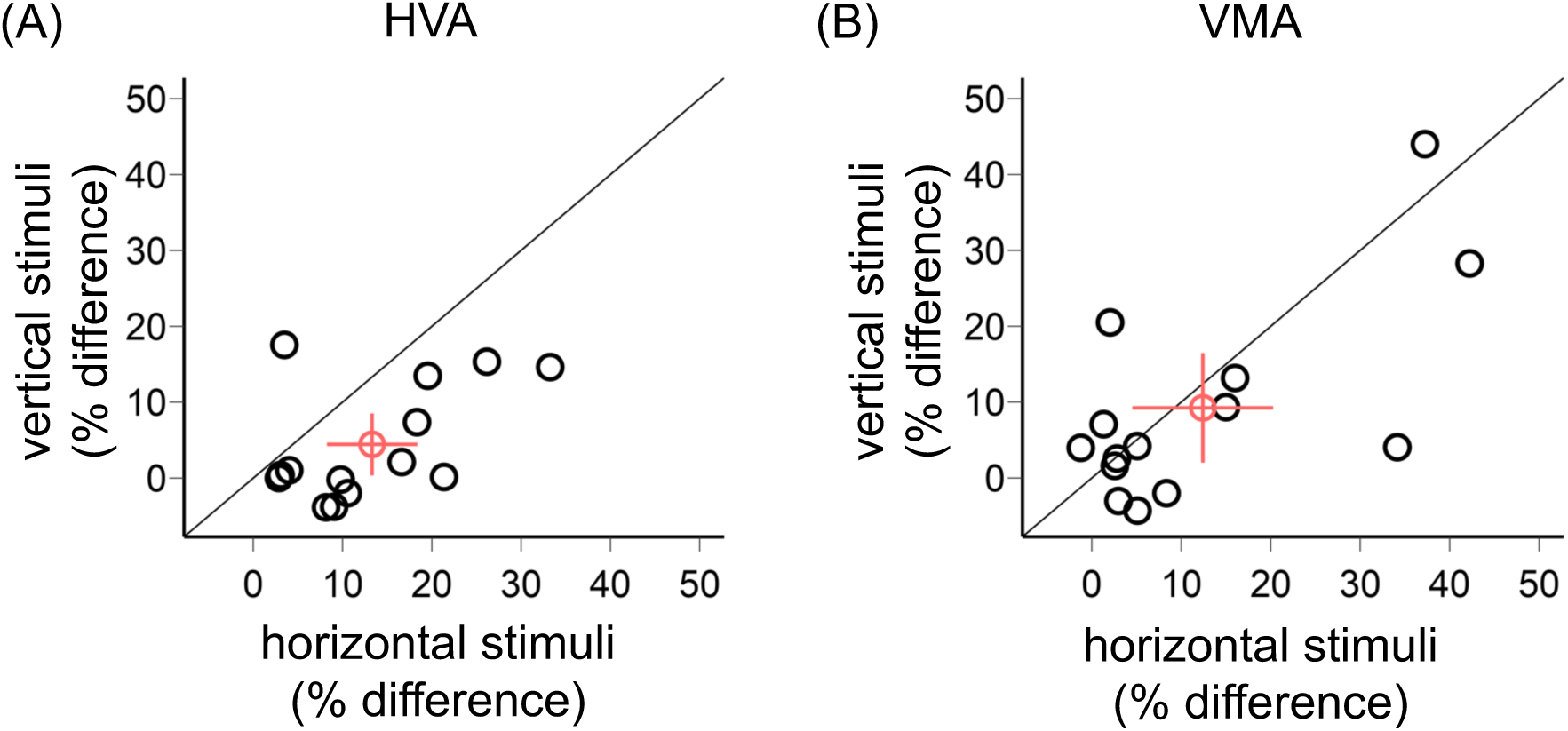
Comparison of HVA (A) and VMA (B) between horizontal and vertical stimuli (% difference in contrast threshold). Each black circle represents the asymmetry for an individual participant; the red circle indicates the mean across participants. Error bars represent ±1 SEM.

We observed no significant correlations between the non-adapted threshold and the adaptation effect for each participant (both Experiments 1 and 2, *ps*>.1), so that the extent of adaptation did not depend on the initial contrast threshold in each condition.

We found a positive correlation between normalized adaptation effects for horizontal and vertical stimuli [*r*=0.46, *p*<.001, **Figure 5A**]. To evaluate the contributions of between-observer and polar-angle variability to these correlations, we regressed out these factors as described in prior research (71). First, we accounted for between-observer variability by subtracting each observer’s average adaptation effect and V1 surface area values across the four polar angle locations. This correlation remained significant after removing between-observer variability [*r*=0.53, *p*<.001, **Figure 5B**]. Additionally, when we removed variability across polar angles by subtracting the average adaptation effect and V1 surface area values for each polar angle across the 13 observers, the correlation still holds [*r*=0.3, *p*=.026, **Figure 5C**]. These findings indicate that the adaptation effect was consistent with both stimulus orientations at the group and the individual levels.

**Figure 5.**
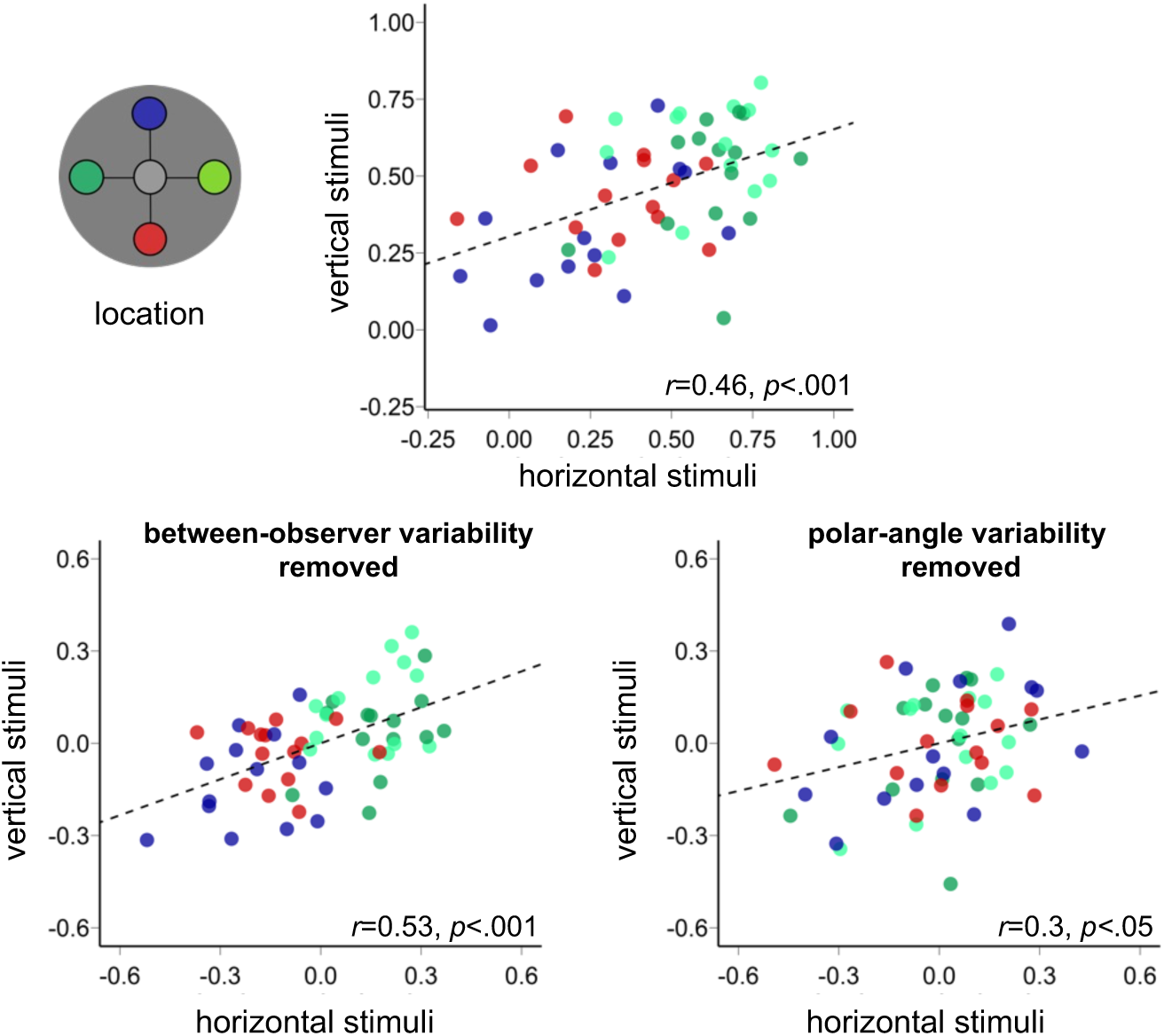
**(A)** Correlation between the normalized adaptation effect ((adapted – non-adapted threshold) / (adapted + non-adapted threshold)) for the horizontal stimulus (x-axis) and the vertical stimulus (y-axis). **(B)** Correlation after removing between-subject variability. (**C**) Correlation after removing polar-angle variability. The dashed black line represents the linear fit to the data points.

In summary, the stronger HVA for horizontal stimuli aligns with a radial bias(106–108). The adaptation effect was stronger at the horizontal than vertical meridian, regardless of the stimulus orientation.

### Linking brain and behavior at perifoveal locations

To test the hypothesis that cortical surface area or pRF size is related to the adaptation effect, we assessed the relation between the normalized adaptation effect and the V1 surface area for 13 out of 14 participants (one participant preferred not to be scanned). Consistent with previous studies(70, 71, 91, 115), V1 surface area was larger along the horizontal than the vertical meridian [*t*(12)=7.51, *p*<.001, *d*=2.08], and along the lower than upper vertical meridian [*t*(12)=2.37, *p*=.035, *d*=0.66, **Figure 6**]. Also consistent with previous studies (70, 71), a correlation indicates that contrast threshold decreases as cortical surface area increases (*r*=-0.35, *p*=.01; **Fig S2, left panel**), this correlation is preserved when we removed between-observer variability (*r*=-0.46, *p*<.001; **Fig S2, middle panel)**, but not when we removed polar-angle variability (*r*=0.05, *p*=.733; **Fig S2, right panel)**. These results highlight the importance of polar angle.

**Figure 6.**
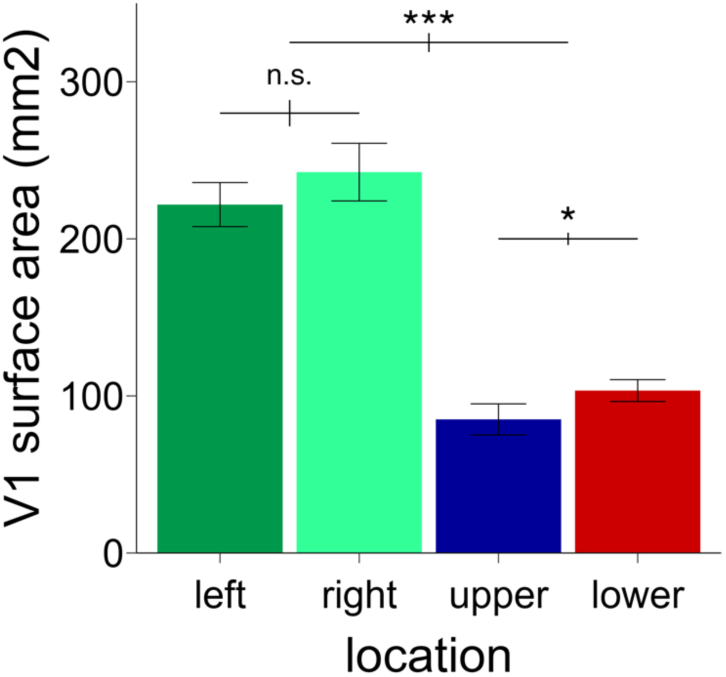
V1 surface area across the four polar angle locations (extended from 4° to 12° eccentricity). Error bars represent ±1 SEM. The error bars above the bar plots indicate ±1 SEM of the difference between conditions \*\*\**p*<.001, \**p*<.05, *n.s. p*>.1.

We observed positive correlations between the normalized adaptation effect and the V1 surface area for both horizontal [*r*=0.56, *p*<.001] (**Figure 7A**) and vertical [*r*=0.37, *p*=.007] (**Figure 7B**) stimuli (top panel). As this correlation relies on the variability across polar angles within the same observers, the data points are not independent. After regressing out the between-observer variability, the positive correlations persisted for both stimulus orientations (horizontal stimulus: *r*=0.73, *p*<.001; vertical stimulus: *r*=0.46, *p*<.001; **Figure 7**, lower left panel). However, when we removed variability across polar angles, the correlations were no longer significant (horizontal stimulus: *r*=-0.003, *p*=.981; vertical stimulus: *r*=0.07, *p*=.632; **Figure 7**, lower right panel). These findings indicate that the observed correlations between the adaptation effect and V1 surface area depend on the polar angle location. Indeed, averaging the V1 surface area and adaptation effect across polar angle locations eliminated the correlations (horizontal stimulus: *r*=-0.19, *p*=.535; vertical stimulus: *r*=0.08, *p*=.793). We observed the same patterns when correlated the non-normalized adaptation effect (computed by subtracting the thresholds) and V1 surface area (Figure S3).

**Figure 7.**
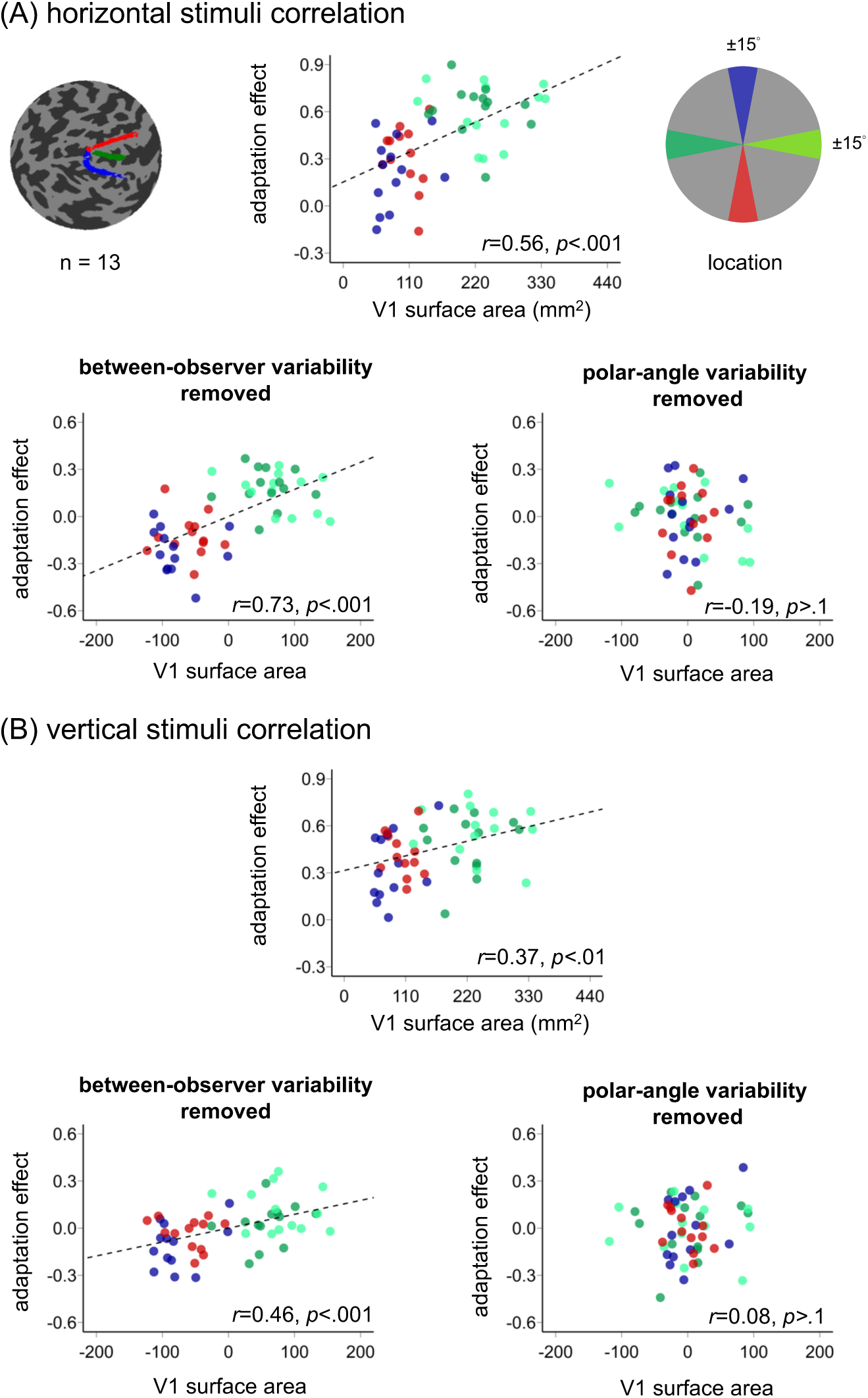
Correlations between the normalized adaptation effect and V1 surface area around polar angle for (**A**) the horizontal stimuli and (**B**) vertical stimuli. Correlations are shown overall (top panels), after removing between-subject variability (lower left panels), and after removing polar-angle variability (lower right panels). The dashed black line represents the linear fit to the data points.

### Experiment 3 – Foveal Location, Horizontal and Vertical Stimuli

The difference of cortical surface area can account for the difference of HVA but not the VMA. To further investigate the relation between V1 cortical surface area and adaptation, we examined the adaptation effect in the fovea and compare it with the one at the perifoveal locations. V1 surface area across eccentricity correlates with various perceptual measures, including perceived object size (65, 66), perceived angular size (67), and acuity (68, 69).

If the visual cortex surface area is the only determining factor of the extent of visual adaptation, the adaptation effect should be stronger at the fovea than the parafovea, and this superiority at fovea should be eliminated by adjusting the target size according to a cortical magnification factor. However, if other factors are also involved in the eccentricity difference, the adaptation effect should differ between the foveal and perifoveal locations.

A two-way ANOVA on contrast thresholds yielded main effects of adaptation [*F*(1,13)=17.69, *p*=.001, *η_p_^2^*=0.58] and stimulus orientation [*F*(1,13)=9.09, *p*=.01, *η_p_^2^*=0.41] but no interaction between them [*F*(1,13)<1]. Contrast thresholds increased after adaptation, and threshold were lower for horizontal than vertical stimuli.

We compared the normalized adaptation effect in the fovea and the perifovea. A three-way ANOVA on adaptation x location x orientation revealed an interaction: *F*(1,13)=5.19, *p*=.04, *η_p_^2^*=0.29, Thus, we conducted separate two-way location x adaptation ANOVAs for horizontal and vertical stimuli.

For the horizontal stimulus, there were main effects of location [*F*(1,13)=43.93, *p*<.001, *η_p_^2^*=0.77] and adaptation [*F*(1,13)=49.25 *p*<.001, *η_p_^2^*=0.79], as well as an interaction [*F*(1,13)=33.72, *p*<.001, *η_p_^2^*=0.72]. The adaptation effect (calculated as the difference between adapted and non-adapted thresholds, as in previous studies (112–114)) was stronger in the perifovea than the fovea [*t*(13)=5.58, p<.001, *d*=1.49]. The same result emerged for the normalized adaptation effect [*t*(13)=7.02, *p*<.001, *d*=1.86] (**Figures 2A, 3C**). There was no correlation between the adaptation effect at the fovea and perifovea, either without normalization (*r*=0.25, *p*=.378) or with normalization (*r*=-0.08, *p*=.777).

For the vertical stimulus, we observed the same patterns. There were main effects of location [*F*(1,13)=44.81, *p*<.001, *η_p_^2^*=0.78] and adaptation [*F*(1,13)=41.39, *p*<.001, *η_p_^2^*=0.76], as well as an interaction [*F*(1,13)=34.79, *p*<.001, *η_p_^2^*=0.73]. Again, adaptation effects were stronger in the perifovea than fovea (**Figures 2B, 3D**) [*t*(13)=6.31, *p*<.001, *d*=1.69], and this was also the case for the normalized adaptation effect [*t*(13)=5.9, *p*<.001, *d*=1.58]. Additionally, there was no correlation between the adaptation effect at fovea and perifovea, either without normalization (*r*=0.28, *p*=.332) or with normalization (*r*=0.31, *p*=.287).

In summary, adaptation effects were consistently stronger in the perifovea than the fovea, irrespective of stimulus orientation.

## Discussion

Here, after confirming performance asymmetries in the non-adapted conditions, we uncovered stronger contrast adaptation effects at the horizontal than the vertical meridian, and in perifoveal than foveal locations, for both horizontal and vertical stimuli. Theses polar angle differences support our horizontal meridian dominance hypothesis: Locations with larger cortical surface areas, which contain more neurons and stronger adaptor representations (96, 97), exhibit a stronger adaptation effect (54, 98–101, 116). Critically, adaptation reduced the HVA, promoting more homogenous perception around the visual field. This differential adaptation effect was mediated by the larger cortical surface area at the horizontal than vertical meridian.

In the non-adapted condition, we observed the typical HVA and VMA. These asymmetries likely arise from both retinal and cortical factors. For example, retinal cone density is higher at the horizontal than vertical meridian (117, 118), midget-RGC density is higher at the lower than upper vertical meridian (118, 119), and V1 cortical surface area is larger for the horizontal than the vertical meridian, and for the lower than upper vertical meridian (**Figure 4**; (63, 70–72, 91). Cortical surface area accounts for more variance in behavioral asymmetries than retinal factors (120). Additionally, factors such as sensory tuning and/or neuronal computations may also contribute to these perceptual asymmetries(3, 92, 121).

This study also revealed that the HVA, but not the VMA, was stronger for horizontal than vertical stimuli– a finding that reflects a radial bias (106–108). Horizontal stimuli favored performance along their radial (horizontal) meridian, whereas vertical stimuli favored performance along their radial (vertical) meridian, thereby respectively potentiating and diminishing the HVA.

The stronger adaptation effect for vertical stimuli (2.5° tilt) than horizontal stimuli (6° tilt) – despite matched performance, so adaptation could have the same opportunity to exert its effect in both experiments– are consistent with orientation dependency in orientation strength: adaptation decreases as the orientation difference between the target and adaptor increases (33, 39, 45, 122–126). However, normalizing the tilt angle (the difference in thresholds between the adapted and non-adapted conditions divided by their sum) eliminated orientation-related differences. Further, the adaptation effects using horizontal and vertical stimuli were positively correlated (**Figure 5A**), regardless of between-subject (**Figure 5B)** or polar-angle variability (**Figure 5C)**. This finding indicates that adaptation effects to two orientations had similar underpinning and were robust and reliable within individuals.

Contrast sensitivity asymmetries with this orientation discrimination task emerge at threshold (6, 10, 88) and at suprathreshold (6, 7, 88) levels. However, neither us nor anyone else has conducted adaptation experiments at suprathreshold levels around polar angle. Thus, whether the present findings can be generalized to higher contrast levels around polar angle remains an open question.

Prolonged exposure to an oriented stimulus causes nearby orientation to appear perceptually shifted away from the adapted orientation (e.g. 127, 128). The tilt angles used in this study were unlikely to trigger a repulsion effect because they are close to the adaptor, within a range reported to yield a slight repulsion (129), and the repulsion is typically shown when the tilt angle is around 10-20 deg (130). Had there been a repulsion effect, we would have predicted a lower threshold in the adapted than non-adapted condition; we observed the opposite. In any case, any repulsion effect would have occurred to the same extent at all tested locations. Thus, a repulsion effect could not explain the differential adaptation effect across meridians. Adaptation reduced contrast sensitivity more along the horizontal than the vertical meridian, regardless of stimulus orientation, even after normalizing the adaptation effect.

Could natural statistics play a role on this differential effect? It has been proposed that asymmetries about stimulus properties, color and orientation, are related to spatial statistics of scenes. For example, the oblique effect, better performance for horizontal and vertical than oblique lines, is related to natural statistics of the visual environment (131–135). Likewise, asymmetries in color space have been related to scene statistics (136). However, as far as we know, there is no existing evidence showing that natural statistics can explain either the horizontal-vertical meridian anisotropy (HVA) or the vertical-meridian asymmetry (VMA). Future studies should explore the contribution of scene statistics to these location asymmetries.

Why was the adaptation effect stronger along the horizontal meridian? Among the few contrast adaptation studies that specified tested locations, adaptation was either assessed throughout the entire visual field (54, 122, 126, 137) or exclusively along the horizontal meridian (55–58). To elucidate a possible mechanism underlying the stronger adaptation effect at the horizontal meridian, we consider the following points: (1) Neurostimulation studies have revealed that V1 plays a causal role in adaptation (36, 42). (2) A positive correlation exists between contrast sensitivity and V1 surface area (71). (3) V1 surface area is also positively correlated with the adaptation effect (**Figure 7**, top panel). This correlation persists after controlling for individual differences (**Figure 7**, lower left panel), but not after accounting for polar angle variability (**Figure 7**, lower right panel). (4) Neuronal density is uniform across the visual field (94, 95). Taken together, these findings suggest that the larger surface area and greater number of neurons at the horizontal than the vertical meridian contribute to the stronger adaptation effect at the horizontal meridian.

These more pronounced adaptation effect at the horizontal than vertical meridian is also consistent with the idea that adaptation helps manage limited bioenergetic resources, as there is more expenditure for the larger cortical areas corresponding to the horizontal than vertical meridian, and with results showing that adaptation increases with the strength of the response to the adaptor (102–105).

The surface area explanation, however, does not align well with the similar adaptation effects observed at the lower and upper vertical meridians. The surface area explanation would have predicted a stronger adaptation effect at the lower than upper vertical meridian. Thus, whereas cortical surface area can explain the decline in contrast sensitivity with increasing eccentricity, it does not fully account for polar angle differences in performance (3, 138). Together, these findings suggest that additional factors beyond surface area may contribute to the observed differential adaptation effects.

Moreover, were V1 surface area the sole factor underlying the extent of adaptation, the fovea would be expected to exhibit the strongest effect. However, in this study, even when the foveal stimulus was equated for cortical representation size (96), adaptation was weaker in the fovea than in the perifovea for both stimulus orientations. These findings are not in line with results showing that adaptation increases with the strength of the response to the adaptor (102–105). These findings are consistent with some adaptation studies (58, 59), but differ from others reporting similar adaptation effects between the fovea and periphery (40, 56). Furthermore, no correlation was observed between adaptation effects in the periphery and fovea.

Could orientation tuning play a role? The half-bandwidth of orientation selectivity is approximately 3-9°. In the instances in which location has been specified, these estimates are typically for foveal locations (124, 125) or for a wide eccentricity range (126). One orientation detection study showed that, for equated performance, orientation bandwidth is broader along the horizontal than the vertical meridian but similar between the upper and lower vertical meridian, demonstrating a horizontal-vertical asymmetry but not a vertical meridian asymmetry (121). Future studies could examine whether orientation tuning for adaptation varies across different meridians.

These results support the idea that foveal and peripheral vision are optimized for distinct perceptual processes (1, 2, 139). The observed differential adaptation effects are likely mediated by qualitative rather than quantitative differences in processing (58). The stronger perifoveal adaptation effect in humans is consistent with macaques studies, which show that contrast adaptation is more pronounced in the retinal and geniculate cells of the peripheral magnocellular pathway than in the more foveally located parvocellular pathway (50, 140).

Like adaptation, covert attention, the selective processing of information (2, 6, 80, 84, 86–89, 141–145), also helps manage limited resources (33–36), and these processes interact in the early visual cortex (36). However, opposite to the effect of adaptation, exogenous/involuntary (6, 7, 86, 87) or endogenous/voluntary (35, 88, 89) covert spatial(80, 146, 147) and temporal (148–150) attention enhance contrast sensitivity at the attended location. Whereas adaptation reduces the HVA, spatial (6, 7, 86–89) and temporal (149) attention enhancements are consistent around polar angle. Thus, covert attention neither exacerbates nor alleviates the HVA or VMA.

Presaccadic attention, which enhances contrast sensitivity at the target location immediately before saccade onset, also has different effects: It enhances contrast sensitivity more along the horizontal than the vertical meridian, and least at the upper vertical meridian (151–153). Consequently, presaccadic attention can amplify polar angle asymmetries. Interestingly, individual presaccadic attention benefits negatively correlate V1 surface area at the upper vertical meridian, suggesting that presaccadic attention helps compensate for the reduced cortical surface area and neuronal count at that location (152). Similarly, the weaker adaptation effect along the vertical meridian in the present study may reflect its smaller cortical surface area and fewer neurons available for adaptation suppression.

In conclusion, this study reveals that contrast adaptation is stronger along the horizontal than the vertical meridian and in the periphery than the fovea, regardless of the adaptor and target orientation. Thus, by mitigating the HVA, adaptation contributes to reducing bioenergetic expenditure as well as inherent physiological asymmetries rendering a more uniform visual perception around the visual field. Moreover, consistent with the critical role of V1 plays in adaptation (36, 42), cortical V1 surface area mediates the differential adaptation effects observed between the horizontal and vertical meridians.

## Materials and Methods

### Participants

Fourteen adults (7 females, age range: 22-35 years old), including author HHL, participated in all three experiments. All of them had normal or corrected-to-normal vision. Sample size was based on previous studies on adaptation (36), with an effect size of *d*=1.3, and on performance fields (151), with an effect size of *d*=1.66 for performance in the neutral trials (without attentional manipulation). According to G*Power 3.0 (154), we would need 9 participants for adaptation and 7 participants for performance fields to reach a power=0.9. We also estimated the required sample size for the interaction between adaptation and location, which enable us to assess performance fields, based on a presaccadic attention and performance fields study (151), as attention and adaptation both affect contrast sensitivity (155), albeit in different directions (35, 36). Bootstrapping the observers’ data from that study with 10,000 iterations showed that we would need 12 participants to reach power=0.9 for the interaction analysis. The number of participants here is similar to or higher than in previous adaptation studies (e.g. 51, 59, 144, 156, 157-160). The Institutional Review Board at New York University approved the experimental procedures, and all participants provided informed consent before they started the experiment.

### Apparatus

Participants were in a dimly lit, sound-attenuated room, with their head placed on a chinrest 57 cm away from the monitor. All stimuli were generated using MATLAB (MathWorks, MA, USA) and the Psychophysics Toolbox (161, 162) on a gamma-corrected 20-inch ViewSonic G220fb CRT monitor with a spatial resolution of 1,280 x 960 pixels and a refresh rate of 100 Hz. To ensure fixation, participants’ eye movements were recorded using EYELINK 1000 (SR Research, Osgoode, Ontario, Canada) with a sample rate of 1,000 Hz.

### Stimuli

In Experiments 1 and 2, the target Gabor (diameter = 4°, 5 cpd, 1.25° full-width at half maximum) was presented on the left, right, upper and lower cardinal meridian locations (8° from the center to center). There were four placeholders (length = 0.16°, width = 0.06°) 0.5° away from Gabor’s edge. The fixation cross consisted of a plus sign (length = 0.25°; width = 0.06°) at the center of the screen.

In Experiment 3, the fixation was replaced by a white placeholder, which was the same size as the other placeholders. We adjusted the target Gabor size according to the Cortical Magnification Factor (96) averaged from nasal, temporal, superior, and inferior formulas (3, 163, 164), which yielded a 1.03° diameter and presented it at the center (0° eccentricity).

### Experimental design and procedures

**Figure 1** shows the procedure of the task. In the <u>adaptation</u> condition, at the beginning of each block, participants adapted to a 100%-contrast horizontal Gabor patch (5 cpd) flickering at 7.5 Hz in a counterphase manner presented at the target location for 60 seconds. Each trial started with 2s top-up phase to ensure a continuous adaptation effect throughout the block. In the <u>non-adaptation</u> condition, participants maintained fixation at the center for 20s (without Gabor) at the beginning of each block and for 2s at the beginning of each trial.

After the top-up, there was a 300, 600 or 900 ms jitter before a tilted Gabor was presented for 100 ms. The fixation plus-sign turned green as a response cue. Participants had to judge whether the target was tilted clockwise or counterclockwise off horizontal or vertical in Experiments 1 and 2, respectively. In Experiment 3, they responded off horizontal or vertical in different experimental sessions. The tilt angle was 6° away from the horizontal line and 2.5° away from the vertical line. They were based on pilot data to ensure a similar adaptation effect while avoiding floor or ceiling performance and were within the neurons’ tuning width, the same orientation ‘channel’ (165–167).

A feedback tone was presented when participants gave an incorrect response. The target locations were blocked. Participants were asked to respond as accurately as possible while fixating at the center of the screen throughout the trial. A trial would be aborted and repeated at the end of the block if participants’ eyes position deviated ≥1.25° from the center from the onset of the adaptation top-up until the response cue onset. There were 48 trials in each block, 4 blocks (192 trials per location for each adaptation and non-adaptation conditions) were conducted consecutively at each location.

In Experiment 1, participants completed the adaptation and non-adaptation conditions on different days, with a counterbalanced order. In Experiments 2 and 3, participants conducted the non-adapted condition followed by the adapted condition. In both Experiments 1 and 2, the order of the target locations was counterbalanced across participants. In Experiment 3, the two stimulus orientation conditions were conducted on different days (one observer the same day but an hour apart) to eliminate any carry-over effect. All observers participated in a practice session to familiarize themselves with the task procedure.

### Titration procedures

We titrated the contrast threshold of the Gabor separately for each location (central, left, right, upper, lower) and adaptation condition (adaptation, non-adaptation) with an adaptive staircase procedure using the Palamedes toolbox (168), as in previous studies (36, 84, 151, 169). There were 4 independent staircases for each condition, varying Gabor contrast from 2% to 85% to reach ∼75% accuracy for the orientation discrimination task. Each staircase started from 4 different points (85%, 2%, 43.5% the median, and a random point between 2% and 85%) and contained 48 trials. We averaged the last 8 trials to derive the contrast threshold. The few outlier staircases (3.3%), defined as the threshold 0.5 log_10_ away from the mean of other staircases in that condition (151), were excluded from data analysis.

### Psychometric function fitting

We fitted a Weibull function for the accuracy as a function of contrast threshold. For each condition, a logistic function was fit to the data using maximum likelihood estimation using the fmincon function in MATLAB. The results derived from the psychometric function estimation positively correlated (*ps<*.01) with the staircase results in all experiments, verifying our procedure in all conditions.

### Behavioral data analyses

Behavioral data analyses were performed using R (170). In Experiments 1 and 2, a two-way repeated-measures analysis of variance (ANOVA) on contrast threshold was conducted on the factors of location (left, right, upper, lower) and adaptation (adapted, non-adapted) conditions to assess statistical significance. We also compared the thresholds in a 3-way repeated-measures ANOVA on the factors of location (left, right, upper, lower), adaptation (adapted, non-adapted), and stimulus orientation (vertical, horizontal) across Experiments 1 and 2. In Experiment 3, we compared the contrast threshold in the fovea and periphery by pooling the performance across all locations in the periphery.

Repeated-measures ANOVA along with effect size (*η^2^*) were computed in R (170) and used to assess statistical significance. *η_p_^2^* was provided for all *F* tests, where *η_p_^2^*=0.01 indicates small effect, *η_p_^2^*=0.06 indicates a medium effect, and *η_p_^2^*=0.14 indicates a large effect. *Cohen’s d* was also computed for each post-hoc *t*-test, where *d*=0.2 indicates a small effect, *d*=0.5 indicates a medium effect, and *d*=0.8 indicates a large effect (171).

The adaptation effect was quantified as the difference between the adapted and non-adapted threshold. We also quantified the normalized adaptation effect based on [(adapted threshold – non-adapted threshold) / (adapted threshold + non-adapted threshold)], similar to quantification of attentional effects (e.g. 100, 172, 173), which takes into account the baseline difference in the non-adapted condition.

### The pRF analysis and correlation with the adaptation effect

We were able to obtain population receptive fields (pRF;(174) and anatomical data for 13 out of 14 observers from the NYU Retinotopy Database (63). One participant preferred not to be scanned. The pRF stimulus, MRI, and fMRI acquisition parameters and preprocessing, the implementation of the pRF model, and the calculation of V1 surface area were identical to those described in the previous work (3, 63, 152). In brief, we computed the amount of V1 surface area representing the left HM, right HM, upper VM, and lower VM by defining ±15° wedge-ROIs in the visual field (centered along the 4 cardinal locations) using the distance maps, as in previous studies (e.g. 71, 72). The cortical distance maps specify the distance of each vertex from the respective cardinal meridian (in mm), with the distance of the meridian itself set to 0 mm. Each ROI extended from 4° to 12° eccentricity. We did not analyze the cortical surface corresponding to the fovea because noise in the pRF estimates of retinotopic coordinates near the foveal confluence tends to be large (63, 72, 175, 176), and the fixation task covered the central 0.5° of the display during the pRF mapping measurement.

The amount of V1 surface area (in mm^2^) was calculated using the average distance of a pool of vertices whose pRF polar angle coordinates lie near the edge of the 15° boundary in visual space, and we excluded vertices outside 30° away from the wedge-ROI center to preclude the noise. Two researchers (including the first author HHL) independently drew the distance maps for the dorsal and ventral part of V1 by hand using neuropythy (https://github.com/noahbenson/neuropythy;(177). The horizontal distance map was derived by the average of the dorsal and ventral maps. These steps were completed for left and right hemisphere of V1 respectively. We then summed through the vertices that had distance within the mean distance calculated for reach cardinal location. Total V1 surface area was highly consistent between independent delineations by two researchers (*r*=0.99, *p*<.001). We then averaged the calculated V1 surface area between the ROIs drawn by the two researchers, as in a previous study (178), and then conducted correlation analysis to evaluate the relation between V1 surface area and the adaptation effect at the individual level.

## Acknowledgments

This research was supported by NIH NEI R01-EY027401 to M.C. and the Ministry of Education in Taiwan to H.H.L. We thank Rania Ezzo, Marc Himmbelberg, Jan Kurzawski, and David Tu for their help in fMRI data acquisition and analysis. We also thank Carrasco Lab members, especially Marc Himmelberg, Ekin Tünçok and Shutian Xue, for their helpful comments on the manuscript.

## Source Data

Data used in this study are available at https://github.com/CarrascoLab/adaptationPF.

