## Supplementary for "Visual adaptation stronger at horizontal than vertical meridian: Linking performance with V1 cortical surface area"

**This PDF file includes:**

Figures S1-S3

### Supplementary Figure

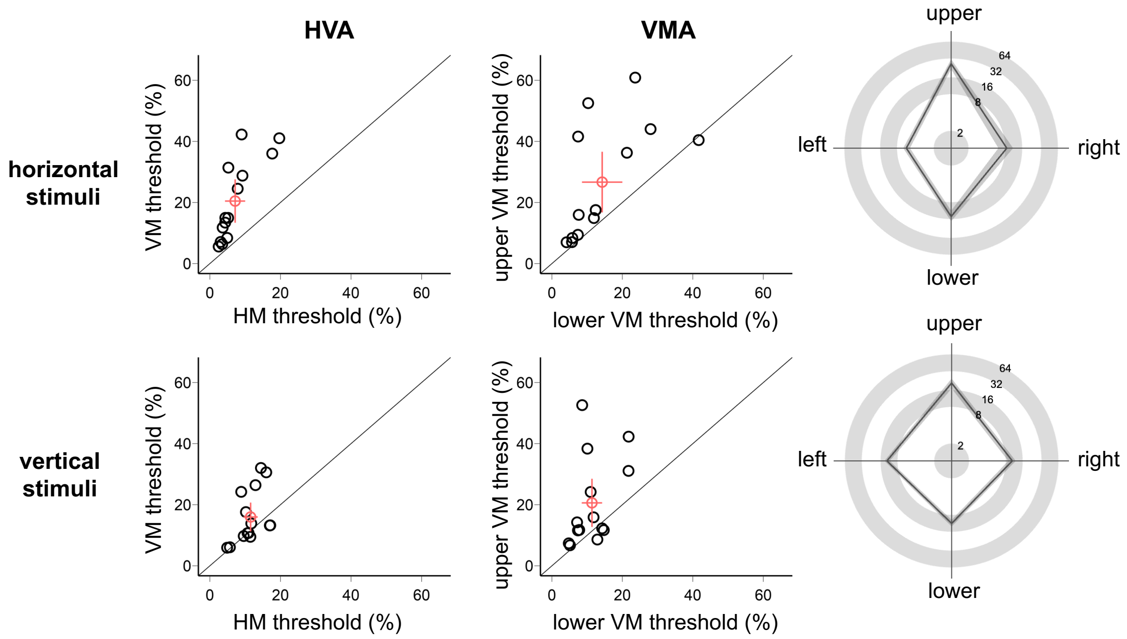


**Fig S1.** Non-adapted thresholds around polar angles for the horizontal stimulus (upper row, Experiment 1) and the vertical stimulus (lower row, Experiment 2). Each dot represents the threshold of each participant. The red dot indicates the mean across participants. Error bars represent ±1 SEM.

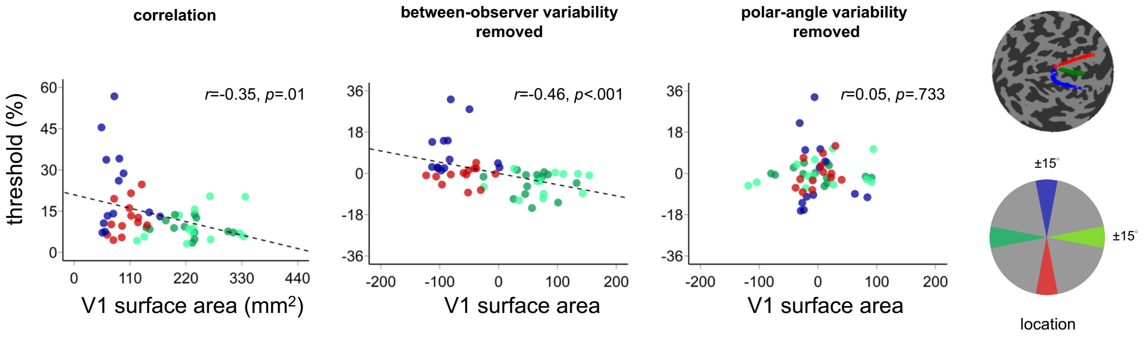


**Fig S2.** Correlations between the averaged contrast thresholds (Experiment 1 and Experiment 2) and V1 surface area. Correlations are shown overall (left panel), after removing between-subject variability (middle panel), and after removing polar-angle variability (right panel). The dashed black line represents the linear fit to the data points.

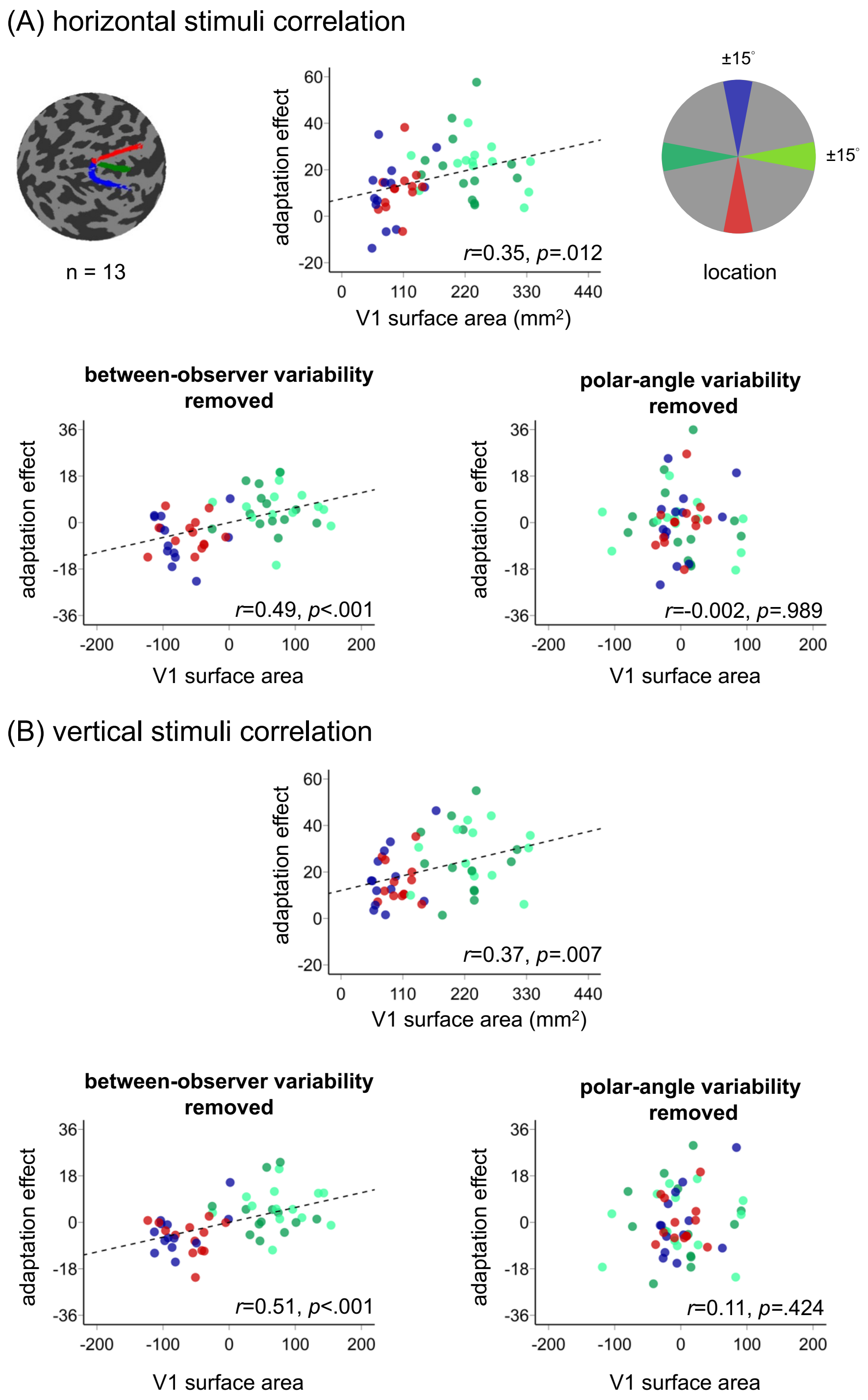


**Fig S3.** Correlations between the non-normalized adaptation effect and V1 surface area around polar angle for (**A**) the horizontal stimuli and (**B**) vertical stimuli. Correlations are shown overall (top panels), after removing between-subject variability (lower left panels), and after removing polar-angle variability (lower right panels). The dashed black line represents the linear fit to the data points.
